# Horizontal Gene Transfer Can Help Maintain the Equilibrium of Microbial Communities

**DOI:** 10.1101/201426

**Authors:** Yuhang Fan, Yandong Xiao, Babak Momeni, Yang-Yu Liu

## Abstract

Horizontal gene transfer and species coexistence are two focal points in the study of microbial communities. The evolutionary advantage of horizontal gene transfer has not been well-understood and is constantly being debated. Here we propose a simple population dynamics model based on the frequency-dependent interactions between different genotypes to evaluate the influence of horizontal gene transfer on microbial communities. We find that both structural stability and robustness of the microbial community are strongly affected by the gene transfer rate and direction. An optimal gene flux can stablize the ecosystem, helping it recover from disturbance and maintain the species coexistence.

## I. INTRODUCTION

Diverse microbial communities, consisting of interacting microorganisms, are essential for their hosts [1–4] or environment [5, 6]. The mechanisms that govern the coexistence of many species in such complex ecosystems are intriguing [7–10]. Another intriguing fact is the presence of horizontal gene transfer (HGT), i.e., the movement of genetic material between microorganisms other than by the transmission of DNA from parent to offspring [11]. Conferring greatly different phenotypes on different strains or species, HGT is central to microbial evolution [12–15]. However, with asexual reproduction being a faster process with lower cost, competition between strains or species can make a beneficial allele spread more efficiently via vertical transfer, rendering a barrier to HGT and contradicting to experimental observations [16]. How HGT can so strongly influence microbial communities still remains an open question [17, 18]. Recently, HGT was observed to play an important role in biofilm formation and public good interactions [19–21], suggesting a possible link between HGT and the stable equilibrium within microbial communities. Furthermore, the boosted HGT was found upon inflammation in the mouse gut microbiota, which can be associated with the dysbiosis of gut microbial community [22], suggesting that HGT could be related to the microbiota dysbiosis. To date, the impact of HGT on the coexistence of different strains within microbial communities has not been fully understood.

Here we propose a simple model to evaluate the impact of HGT on microbial population dynamics. This model can help us understand the evolutionary force that drives HGT and explain how different genotypes can coexist stably in microbial communities. Our results can naturally explain the existence of stable gene flux and reveal how gene transfer rate and direction can affect the microbial system’s structural stability and robustness.

## II. MODEL

Consider a community composed of two genotypes, one containing a cooperative allele and the other containing a cheating allele. The microbes with cooperative alleles will behave cooperatively to benefit both themselves and the whole community, while microbes with the cheating allele will only benefit themselves. We assume that these two alleles can transfer horizontally within the microbial community, making microbes alter their behavior accordingly [23–25]. Without loss of generality, we let *C* be the fraction of cooperators and (1 − *C*) be the fraction of cheaters. The population dynamics of the microbial community can be described by an ordinary differential equation (ODE) [18, 26]:

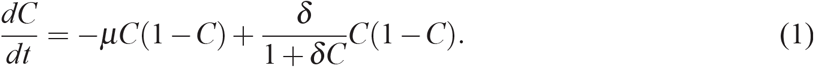

The first term −*μC*(1 − *C*) represents the effect of HGT and the second term *C*(1 − *C*)*δ*/(1 + *δC*) represents natural selection. Here *μ* is the rate of horizontal gene transfer with positive sign representing a strong cheating allele transfer effect. *δ* is the selective coefficient of the cooperators.

In conventional population genetic studies, gene alleles are simply classified as beneficial or harmful without considering the concrete population structures [27]. Therefore a genotype’s fitness is typically constant. However, it has been revealed that the fitness brought by a gene allele can highly depend on the population structure of microbial communities [28, 29]. As a result of complex and nonlinear interactions among different genotypes within the microbial community [30], an initially beneficial gene allele could become harmful for microorganisms carrying it under some specific population structure and vice versa. In other words, we can not simply regard HGT as another way to disperse beneficial allele in evolution. Instead, in our model, we assume that the selective coefficient *δ* is not a constant but a function of the cooperator fraction *C* (Fig. 1). The exact functional form of *δ*(*C*) depends on the properties of the microbial community (see Methods). It turns out that many fundamental properties of the dynamical system (1) can be analytically derived, even without knowing the detailed functional form of *δ*(*C*).

**FIG. 1.**
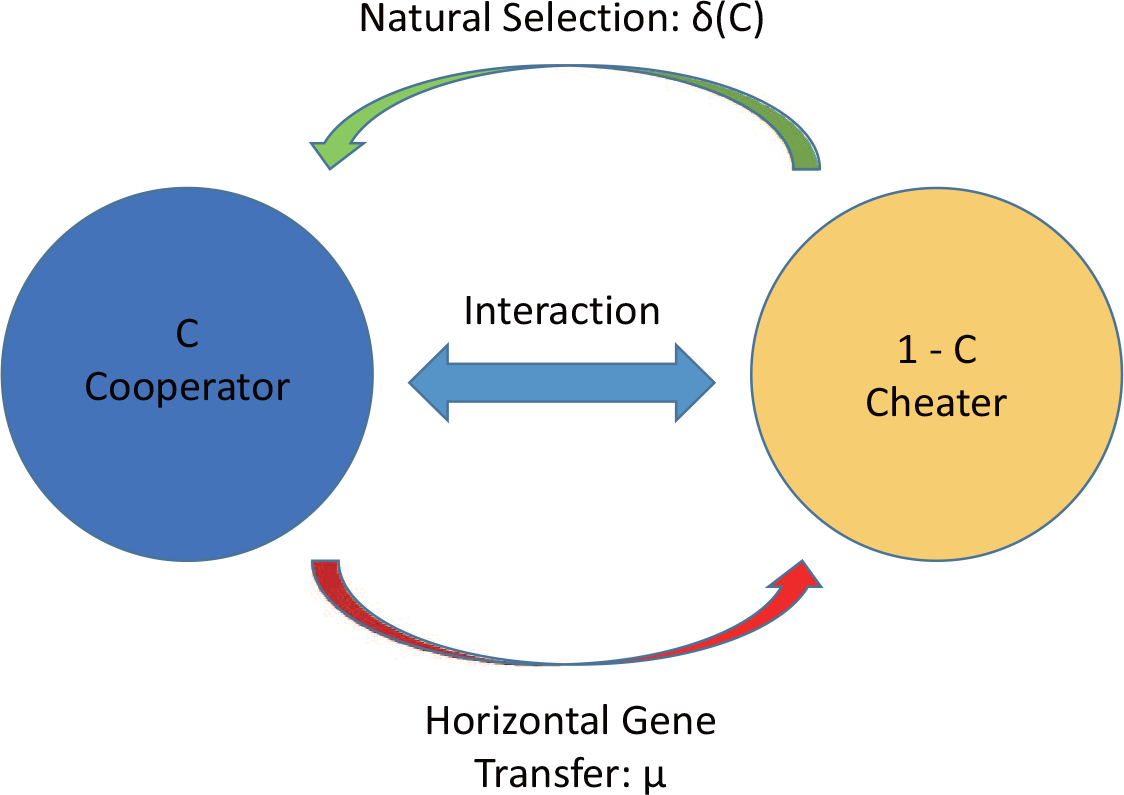
Both natural selection and horizontal gene transfer shape microbial population dynamics. Sub-communities *C* (blue) and 1 − *C* (yellow) represent the fractions of cooperators and cheaters in the microbial community, respectively. Their fitness typically changes according to the interaction among them, rendering a frequency dependent selective coefficient *δ*(*C*). Green and red arrows represent the effect of natural selection and horizontal gene transfer, respectively.

## III. RESULTS

### A. Stability Criteria for Steady States

The population dynamics (1) has two trivial steady states:

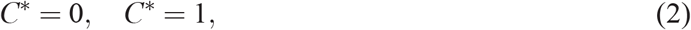

and non-trivial ones (0 < *C*\* < 1) corresponding to the coexistence of cooperators and cheaters,

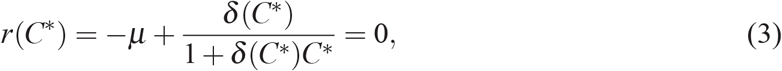

that is, the effects of gene transfer and natural selection balance out. For a steady state *C*\*, the necessary and sufficient condition of its Lyapunov stability is *dC*/*dt* = *r*(*C*)*C*(1 − *C*) < 0 in the right neighborhood of *C*\* and *dC*/*dt* = *r*(*C*)*C*( 1 − *C*) > 0 in the left neighborhood of *C*\*. To prevent our microbial system from evolving to non-coexistent states, we just need to ensure that the two trivial steady states *C*\* = 0 and *C*\* = 1 are unstable. Mathematically, since *C*( 1 − *C*) > 0 always holds on interval (0,1), we have the following stability criteria for the two trivial steady states:

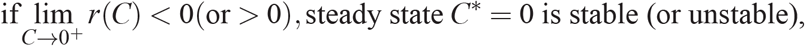

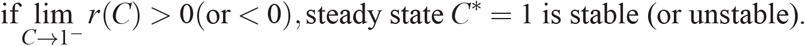

In fact, we can assert from the stability criteria that if neither *C*\* = 0 nor *C*\* = 1 is stable, as a result of the intermediate value theorem, system (1) has at least one non-trivial stable steady state, corresponding to the coexistence of cooperators and cheaters.

### B. Stable gene flux

The presence of stable and long-term gene flux has not been fully understood. Previous studies suggest that only very little gene transfer should occur because the carriers of beneficial allele can rapidly outcompete the other microbes before the beneficial allele has chance to transfer [17, 31], However, in our model the fitness of a genotype is a function of *C*, which can naturally lead to the stable gene transfer. Indeed, if Eq. (3) has a non-trivial solution *C*\* corresponding to a stable steady state on interval (0,1), then a stable positive gene flux rate *l* = |*μC*\*(1 − *C*\*)| exists around the neighborhood of *C*\* (Fig.2b-c). Stable gene flux generally exists in those cases where both natural selection and gene transfer shape population dynamics at equilibrium.

**FIG. 2.**
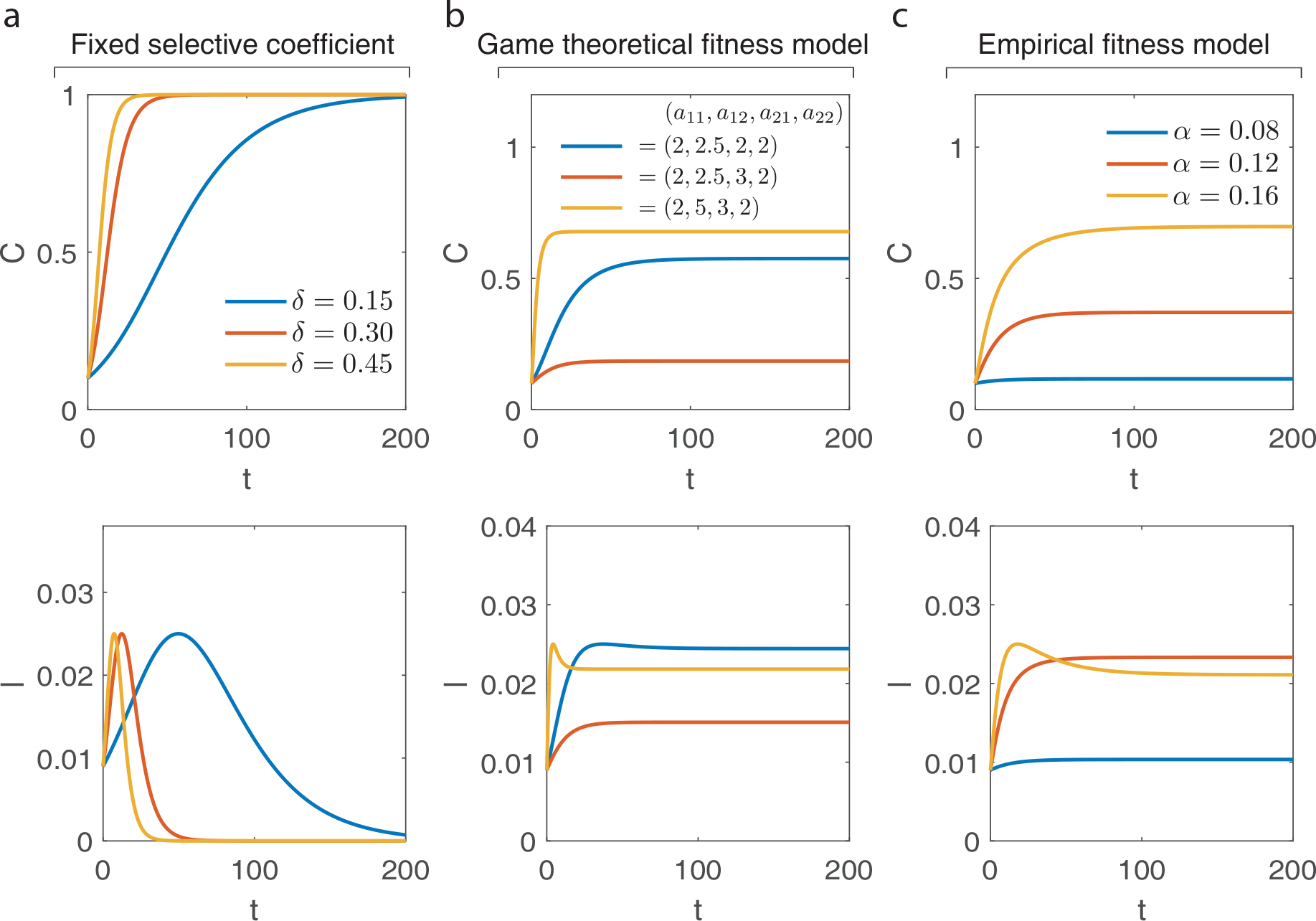
Alterable selective coefficient leads to a stable gene flux. The fraction of cooperators *C* (top) and the gene flux rate *l* = |*μC*( 1 − *C*) | (bottom) are represented as a function of time. **(a)** Fixed selective coefficient (*δ* =constant). Genotype carrying beneficial allele quickly dominates the whole population, leading to zero gene flux. Here we take *μ* = 0.1 in the simulations. **(b)** Game theoretical fitness model. Stable gene flux exists with frequency dependent selective coefficient 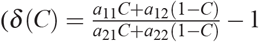, see Methods). Here we take *μ* 0.1 in the simulations, **(c)** Empirical fitness model. Stable gene flux also exists with 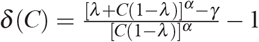 (see Methods). Here we take λ = *δ* = 0.5 and *μ* = 0.1 in the simulations.

By contrast, in conventional theories the gene transfer direction is always from the genotype that has higher fitness to that with lower fitness (see SI Sec. I). Both *μ* and *δ* are constants and have opposite signs [18], Moreover, since very large fitness difference is really rare in practice, one can assume that *δ* > −1 and 1 + *δC* > 0, rendering −*μ* + *δ*/(1+*δC*) ≠ 0. Consequently, the dynamic system (1) has only two trivial steady states: *C*\* = 0 and *C*\* = 1, rendering no coexistence and zero gene flux rate (Fig.2a).

### C. Structural Stability

If starting from any initial state the system can reach a stable steady state with two genotypes coexisting, then we call these two genotypes “globally” coexist. As we discussed above, this occurs if and only if both *C*\* = 0 and *C*\* = 1 are unstable. We call the region of the model parameter space that can guarantee the global coexistence of all genotypes as “flexible coexistent region”. The volume and shape of this region can be used to quantify the *structural stability* of the dynamic system (1).

Structural stability is a classical notion in dynamic systems theory [32]. A system is considered to be structural stable if smooth change in its model parameters will not change its dynamical behavior. Structural stability plays a key role in maintaining ecosystem equilibrium [33–35]. In this work we use the volume and shape of the flexible coexistent region to quantify a microbial system’s structural stability. A larger or more suitable flexible coexistent region will make the microbial system more robust to the environmental disturbance, leading to higher structural stability. We find that HGT can be explained as an efficient way for microbial systems to adjust or even expand their flexible coexistent region to maximize their structural stability in complex environment.

We assume that all the model parameters can affect the selective coefficient of our model, i.e., *δ* = *δ*(*C*,*γ*_1_,…,*γ*_*n*_), where (*γ*_1_,…,*γ*_*n*_) are model parameters. Whether a smooth change of *γ*_*i*_ will change the microbial community’s dynamical behavior determines its structural stability. Based on the stability criteria we derived above, to insure global coexistence, we just need to keep lim_*C*→0+_*r*(*C*) > 0 and lim_*C*→1_- *r*(*C*) < 0. Define 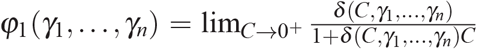 and 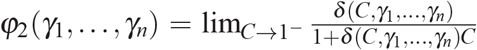. We derive that the flexible coexistent region can be expressed by *φ*_1_(*γ*_1_,…,*γ*_*n*_) > *μ* and *φ*_2_(*γ*_1_,…,*γ*_*n*_) < *μ*. This implies that HGT can naturally shape the flexible coexistent region (Fig.3) and hence impact the system’s structural stability for arbitrary frequency-dependent fitness form *δ*(*C*,*γ*_1_,…,*γ*_*n*_). Furthermore, we can show that the above analysis can be extended to the case of arbitrary number of genotypes (see Methods).

**FIG. 3.**
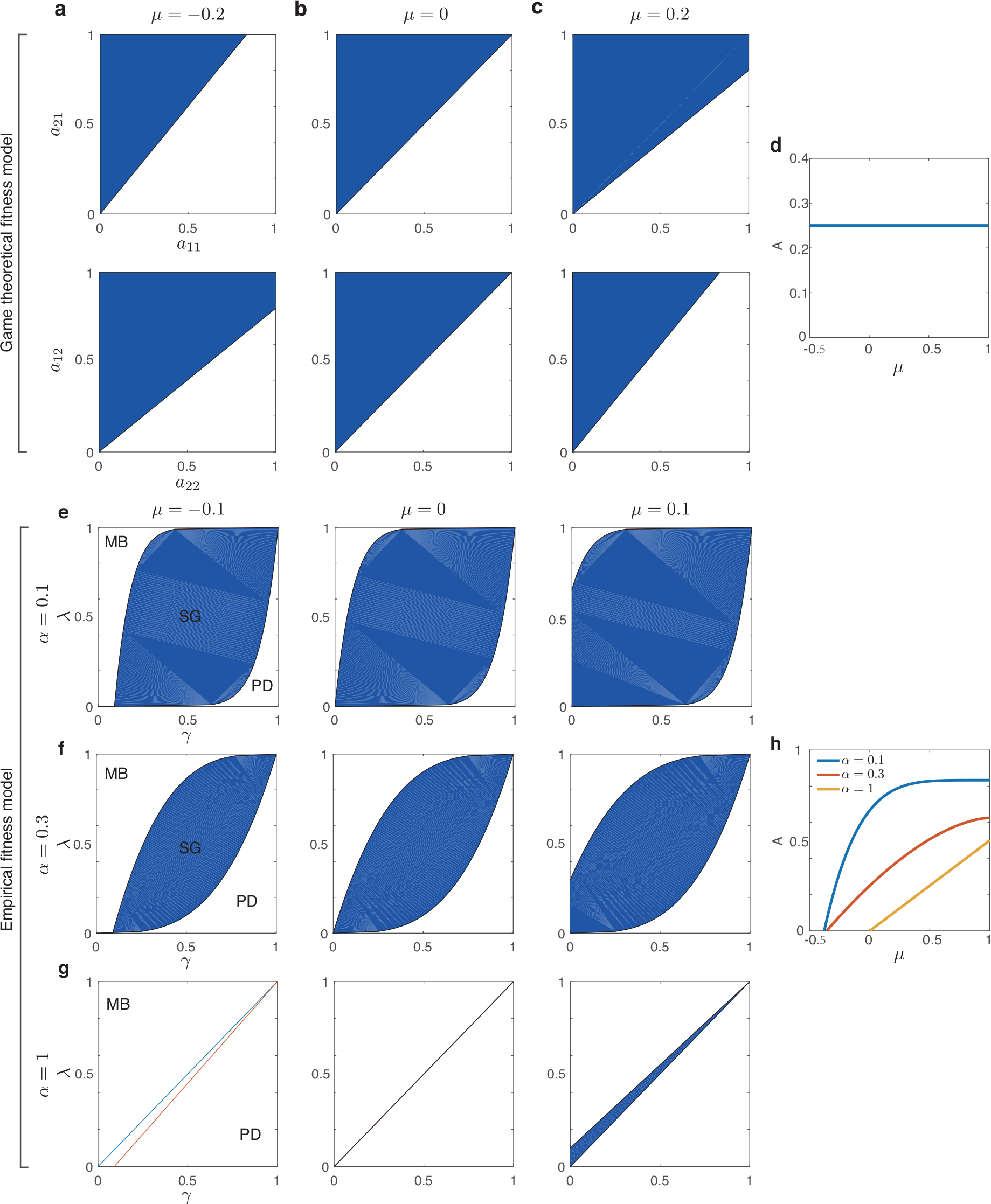
Horizontal gene transfer can adjust the flexible coexistent region of microbial systems. **(a-d)** Game theoretical fitness model. The flexible coexistent regions correspond to the blue areas when the gene transfer rate *μ* = ‒0.20 (a), 0 (b) and 0.2 (c). It can be seen that a positive gene transfer rate expands the flexible region spanned by an *a*_11_ and *a*_21_ but reduces the flexible region spanned by *a*_22_ and *a*_12_. The volume of the flexible coexistent region (*A*, see Methods) spanned by model parameters (*a*_11_, *a*_21_, *a*_21_, *a*_22_) is plotted as a function of the gene transfer rate *μ* in (d). **(e-h)** Empirical fitness model. The flexible coexistent region for the case of *α* = 0.1 (e), 0.3 (f) and 1 (g) with *μ* = ‒0.1, *μ* = 0 and *μ* = 0.1, respectively. Note that low efficiency (*λ*) and high cost (*δ*) lead to the prisoner’s dilemma (PD) regime, while high efficiency and low cost lead to the mutually beneficial (MB) regime. Both regimes have no coexistence. Only in the snowdrift game (SG) regime can the two genotypes coexist. The volume of the flexible coexistent region is plotted as a function of the gene transfer rate *μ* for different *α* in (h). It can be seen that a higher positive gene transfer rate leads to larger flexible coexistent region, thus enhancing the system’s structural ability.

Define *A* to be the volume of flexible coexistent region determined above. The derivation of *A* about HGT rate *μ* can be calculated by the following multiple integral of the first kind:

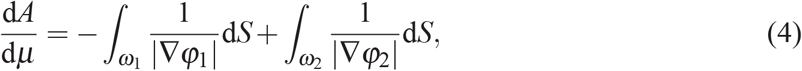

where integral regions *ω*_1_ and *ω*_2_ are boundaries of flexible coexistent region shaped by *φ*_1_(*γ*_1_,…,*γ*_*n*_) > *μ* and *φ*_2_(*γ*_1_,…,*γ*_*n*_) < *μ*, respectively. Intuitively, a higher (lower) HGT rate can reduce (expand) the parameter region where cooperators dominate and expand (reduce) the parameter region where cheaters dominate. Therefore the composite effect of the change of HGT rate is the difference of above two effects of cooperators and cheaters (corresponding to the two multiple integral terms in Eq. (4)). As an extended flexible coexistent region corresponds to higher structural stability, we can evaluate the impact of HGT on a microbial community by considering the sign of d*A*/d*μ*: a positive (or negative) d*A*/d*μ* suggests a positive (or negative) effect of higher HGT rate on the microbial community’s structural stability, respectively.

We validate Eq. (4) for two concrete models: a game theoretical one and an empirical one (see Methods). Interestingly, we find that for the game theoretical fitness model, the volume of the flexible coexistent region spanned by the system parameters *a*_11_, *a*_12_, *a*_21_ and *a*_22_ is invariant with respect to the gene transfer rate *μ*, though its shape does vary over system parameters (Fig.3a-d). For the empirical fitness model (see Methods), increasing *μ* leads to a larger *A* and hence stronger structural stability (Fig.3e-g).

### D. Robustness

In the above analysis of structural stability, we focus on the case of global coexistence, in which microbial system will converge to coexistence of multiple genotypes from any initial state. If Eq. (3) has only one solution in (0,1), then the system will reach a steady state that is independent of the initial condition [36] (Fig.4b-e). However, in practice, ecosystems may have multiple non-trivial steady states. Whether an ecosystem will converge to a particular stable coexistence equilibrium may highly depend on the initial state and the level of environmental disturbances. Indeed, it has been observed that the dynamics of microbial systems often subjects to bifurcation [37–39], that is, if the population of one genotype drops below or over some threshold then the system of coexisting multiple genotypes will collapse and only one genotype will dominate (Fig.4a).

**FIG. 4.**
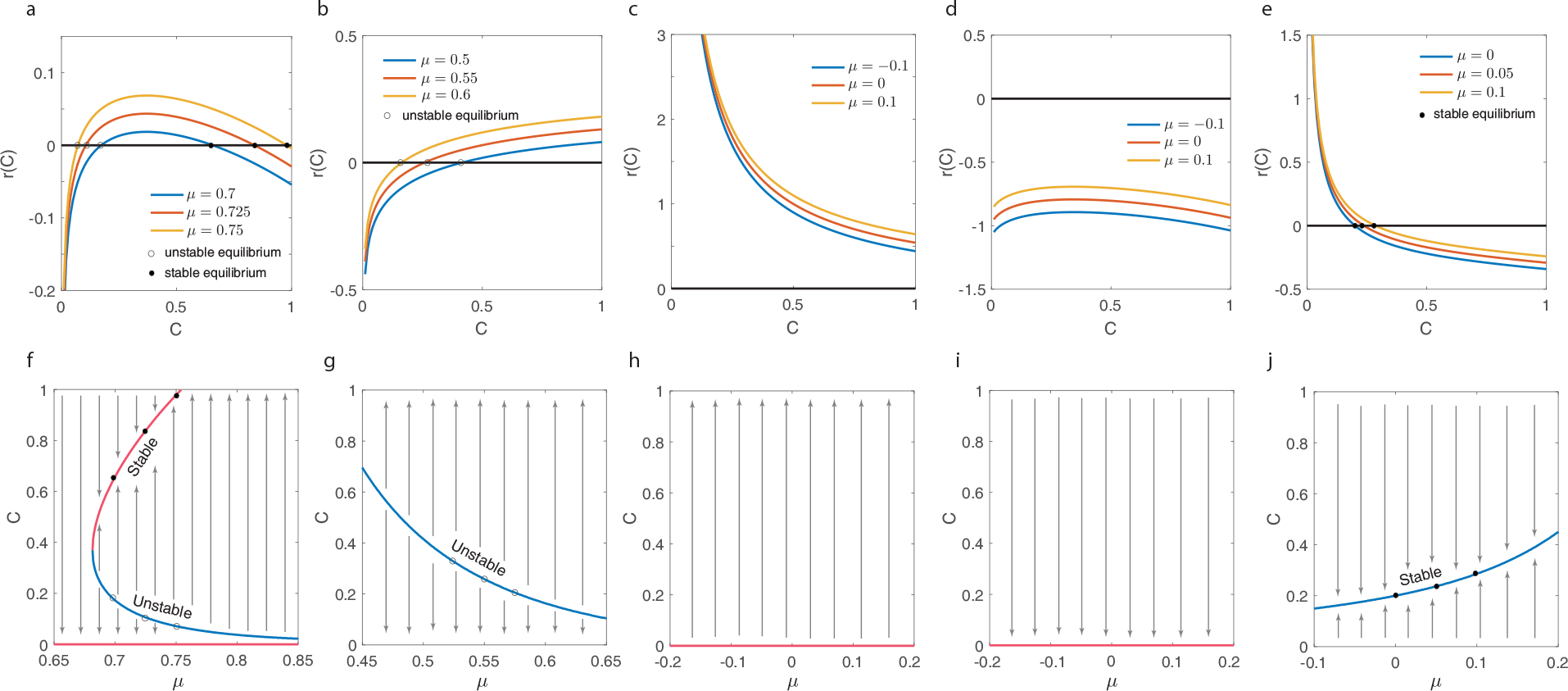
The impact of horizontal gene transfer on the robustness of microbial systems with an empirical fitness model. **(a-e)** The solutions of Eq. (3) correspond to the intersection points with x-axis. The solid circles represent the stable equilibria while the hollow circles mean the unstable steady states, i.e., threshold of according dynamic system. **(f-j)** With the different model parameters (*α, δ, λ*), the selection coefficient *δ*(*C*) (see Eq. (22) in Methods) can lead to various scenarios on the non-trivial equilibria. (a, f) Partial Coexistence. Eq. (3) has two nontrivial solutions corresponding to one stable and one unstable equilibrium, respectively. Whether the microbial system will coverage to a stable coexistent equilibrium highly depends on the initial condition. (b, g) Bi-stable. Only one nontrivial equilibrium exists, but it is unstable. The system will converge to a trivial equilibrium. Whether *C*\* = 0 or 1 depends on the initial condition. (c, d, h, i) One species dominates. No coexistent equilibrium exists. The system will converge to a trivial equilibrium, regardless of the initial condition. (e, j) Global coexistence. Any initial condition will lead to the only non-trivial equilibrium. Hence the two genotypes always coexist. Parameters used for different cases are: (*α, δ, λ*)=(0.765,0.799,0.744) (a, f); (0.69,0.46,0.32) (b, g); (0.55,0.1,0.8) (c, h); (0.3,0.5,0.1) (d, i); (0.5,0.5,0.55) (e, j).

For those systems, we can define robustness as the maximal degree of perturbation such systems can sustain around a stable coexistent equilibrium. A wider range of initial states from which microbial system can evolve to this coexistent equilibrium means a more robust microbial system. Mathematically, if Eq. (3) has multiple solutions on interval (0,1), then we can evaluate the robustness of a stable equilibrium by its minimum distance to its neighbouring unstable equilibrium. From Eq. (3) we know that change of gene transfer rate *μ* can adjust the distance between solutions of *r*(*C*) = 0 (Fig.4f, Fig.S1). Hence HGT is naturally linked to the robustness of such microbial systems. For different dynamic systems and system parameters, the optimal direction and rate of gene transfer that maximize the ecosystem utility could be system-dependent.

## IV. DISCUSSION

Our analysis reveals the key role of HGT in maintaining a microbial system’s equilibrium. By introducing the frequency-dependent fitness and discussing the mutation-selection balance, we offer a simple explanation for the presence of stable gene flux. We quantify the impact of HGT on shaping the flexible coexistence region, and reveal how HGT can influence the structural stability and robustness of microbial communities. Moreover, we validate our scheme for two concrete fitness models where global coexistence can exist. Our analysis can be easily extended to the case of multiple genotypes, suggesting that HGT is an indispensable factor when analyzing microbial population dynamics.

Recently, a theoretical model is proposed to explain the horizontal sweeps via migration among microbial communities [18]. We show that our conclusion still holds under the effect of migration (see SI Sec. III). Note that in our model, for the sake of simplicity, we neglect the cost caused by gene transfer, and the gene transfer rate has no direct impact on organisms’ fitness. However, since it requires cells to produce gene carriers such as plasmid to make HGT happen in practice [40, 41], a higher gene transfer rate can lead to higher cost for microbes which transfer their genes to others. Therefore their fitness could be associated with the gene transfer rate *μ*. We develop an extended model to evaluate this case (see SI Sec. IV). In this extended model, the impact of HGT on microbial system still exists, and adjusting *μ* can potentially lead to an improvement of microbial system’s structural stability and robustness to environmental perturbation. Furthermore, we also develop an individual-based model to confirm our prediction of the impact of horizontal gene transfer on structural stability (see SI Sec. V).

In this work we demonstrate the key link between structural stability, robustness and HGT for a well-mixed and simplified microbial system. The systematic effect of HGT on more complicated and real microbial systems with spatial pattern and various species remains to be studied. We anticipate that introducing the effect of HGT to various processes of microbial systems, especially those associated with coexistence could help us better understand many intriguing phenomena observed in microbial systems.

## V. METHODS

### A. General structural stability model for muti-genotypes microbial system

For a general microbial system with *m* genotypes, we can denote their fitness as a function of population fraction 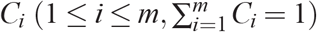, model parameters *γ_j_* (1 ≤ *j* ≤ *n*) and gene transfer rate matrix *μ*, i.e, *f_i_*(*C*_1_,…, *C_m_*,*γ*_1_,…,*γ_n_*, *μ*), 1 ≤ *i* ≤ *m*. Therefore we have that the selective coefficient of genotype *i*

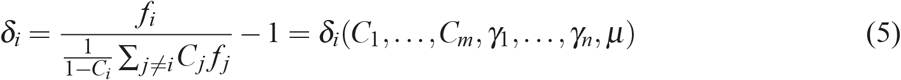

as a function of population fraction and model parameters (see SI Sec. II). Denote the rate of HGT from genotype *i* to genotype *j* is *μ*_*ij*_. Then we have the following dynamic system

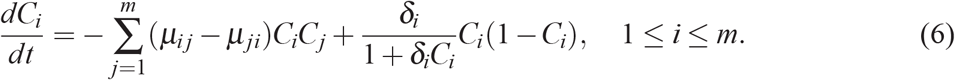

Conditions

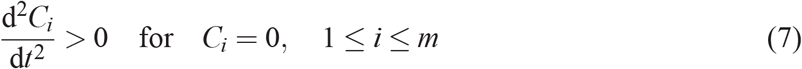

and

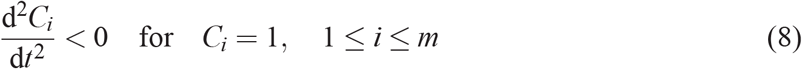

determine the coexistent region of parameter space, which can be expressed as

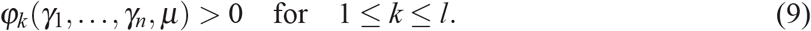

Therefore the flexible coexistent region, i.e., the structural stability of a microbial system is associated with HGT rate *μ*.

### B. Game theoretical fitness model

The concrete functional forms of the selective coefficient, *δ*(*C*), are various for different microbial communities. We first consider a classic game theoretical fitness model with two strategies cooperating and cheating in a microbial community [42, 43]. The payoff matrix (*a_ij_*) for this game (where *a_ij_* represents the payoff of the microbes with strategy *i* when encountering microbes with strategy *j*) is given by

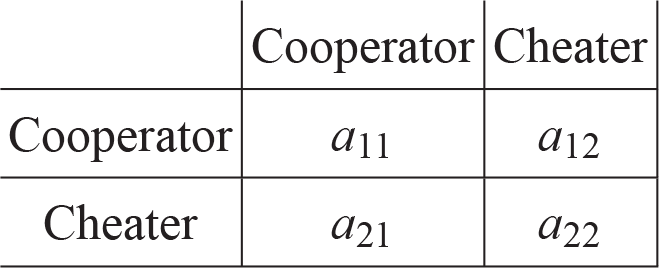

|  | Cooperator | Cheater |
| --- | --- | --- |
| Cooperator | $a_{11}$ | $a_{12}$ |
| Cheater | $a_{21}$ | $a_{22}$ |

For the sake of normalization, we assume that 0 < *a_ij_* < 1 for *i, j* = 1, 2. The fitness of cooperators (*f_C_*) and cheaters (*f_N_*) are given by

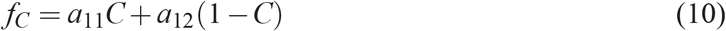

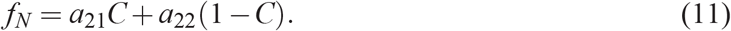

Then the selective coefficient *δ* can be calculated as follows [27]

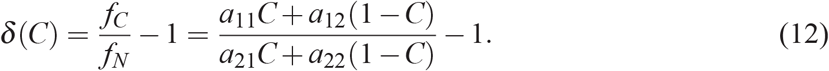

For the case of evolutionary matrix game, we regard elements *a_ij_* in payoff matrix as model parameters. Since

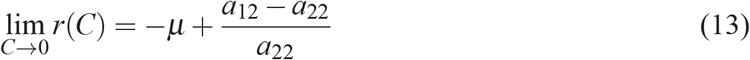

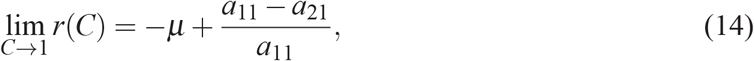

according to our inference mentioned above, we have the flexible coexistent region as

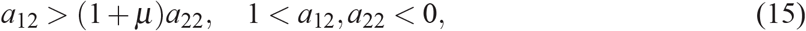

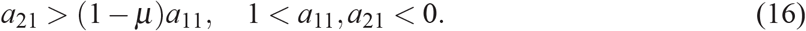

It can be seen that *μ* has a clear impact on the flexible coexistent region where the two genotypes can stably coexist. A positive *μ* will expand the flexible region corresponding to Eq. (16) but reduce the flexible region corresponding Eq. (15) (Fig.3).

Interestingly, we find that HGT does not change the total volume of the flexible coexistent region for game theoretical fitness model. From Eq. (15) and (16) we can calculate the volume of the flexible coexistent region’s projection on the subspace spanned by *a*_11_, *a*_21_ and *a*_12_, *a*_22_, respectively:

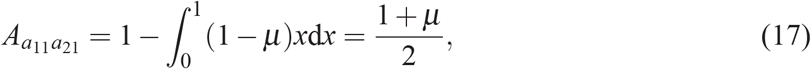

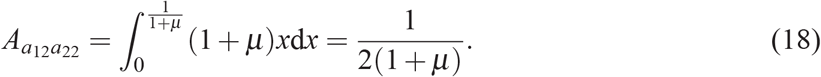

Without loss of generality, here we suppose that *μ* > 0. For *μ* ≤ 0, a similar derivation can be done. Since two conditions shaping the flexible coexistent region are independent, we can get that the volume of flexible coexistent region in the parameter space as

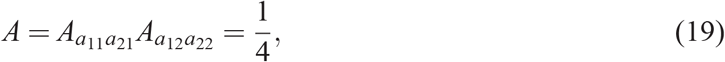

which is independent of *μ*.

We can also calculate d*A*/d*μ* for *μ* > 0 according to Eq. (4):

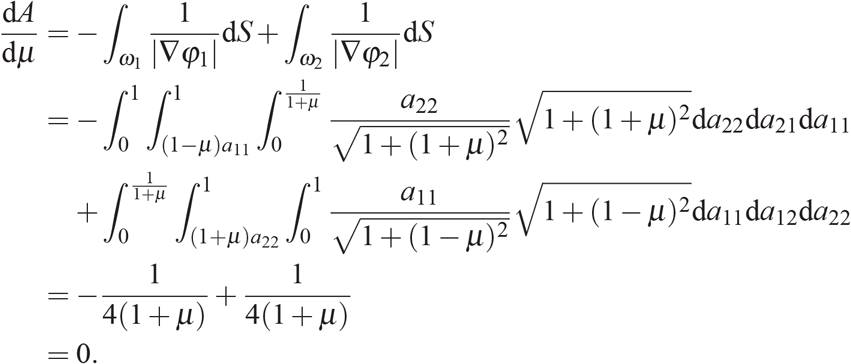

That is, the positive effect and negative effect of HGT cancel out (and the same result also holds for *μ* < 0).

### C. Empirical fitness model

Recently, a snowdrift game dynamics was proposed to explain experimental observation in yeast microbial community [28]. The following functional forms of fitness *f_C_* and *f_N_* were used to fit the experimental observation [28]:

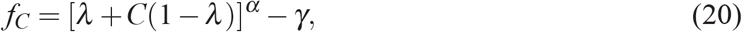

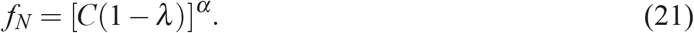

Here *γ* is the cost of cooperation and *λ* is the efficiency of generating total benefits, *α* is a model parameter. With suitable parameter choice, this is a snowdrift game (SG) dynamics. Note that the SG game can result into the coexistence of cooperators and cheaters, which is impossible for the prisoner’s dilemma (PD) or the mutually beneficial (MB) game, where one strategy (genotype) dominates. Selective coefficient *δ* can be then calculated as follows:

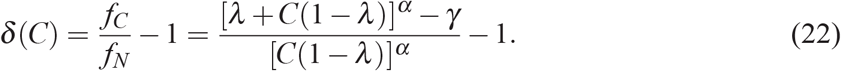

To guarantee the global coexistence of two genotypes, we just need to guarantee that *C*\* = 0 and *C*\* = 1 are unstable steady states. For this empirical fitness model (22) we have

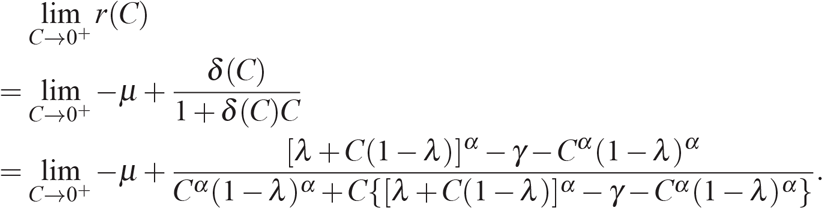

The limitation tends to infinity, which has the same sign as λ^α^ ‒ δ. λ^α^ ‒ δ = 0 is a zero measure on the plane so that we can neglect this case. Moreover, we have

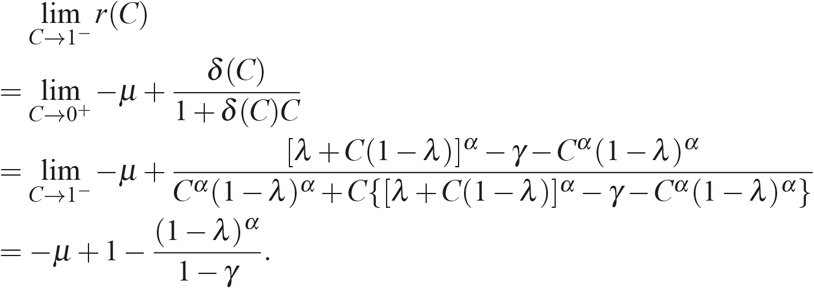

Therefore for this empirical fitness model, with a given *α*, there is a flexible region in the parameter space spanned by *λ* and *γ* where two genotypes can coexist stably with any initial condition. The coexistent region is described as below:

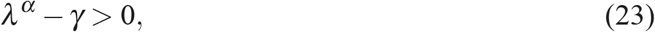

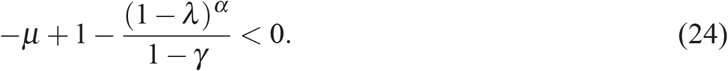

Due to the singularity of function *r*(*C*) near zero, the gene transfer rate *μ* has no impact on the first condition. But clearly *μ* appears in the second condition and a higher positive HGT rate *μ* can lead to a larger flexible parameter region of coexistence, corresponding a higher structural stability. For this case, we have

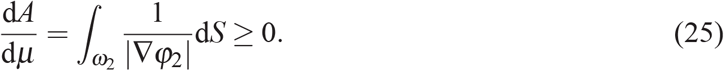

As what we describe above, for comparatively simple case when Eq. (3) only has at most one solution on (0,1), we understand the corresponding ecosystem dynamics very well (Fig.4). However, if Eq. (3) has more than one solution on the interval (0,1), then the robustness of microbial system may be limited.

## Author Contribution

Y.-Y.L. conceived and designed the project. Y.F. performed all the analytical calculations. Y.F. and Y.X. performed numerical simulations. All authors analyzed the results. Y.F. and Y.-Y.L. wrote the manuscript. B.M. and Y.X. edited the manuscript.

## Acknowledgements

This work was partly supported by the John Templeton Foundation under Award No. 51977. Y. F. is sponsored by Undergraduate Overseas Research Fellowship of Cho Kochen Honors College, Zhejiang University.

